# DC/L-SIGN recognition of spike glycoprotein promotes SARS-CoV-2 trans-infection and can be inhibited by a glycomimetic antagonist

**DOI:** 10.1101/2020.08.09.242917

**Authors:** Michel Thépaut, Joanna Luczkowiak, Corinne Vivès, Nuria Labiod, Isabelle Bally, Fátima Lasala, Yasmina Grimoire, Daphna Fenel, Sara Sattin, Nicole Thielens, Guy Schoehn, Anna Bernardi, Rafael Delgado, Franck Fieschi

## Abstract

The efficient spread of SARS-CoV-2 resulted in a pandemic that is unique in modern history. Despite early identification of ACE2 as the receptor for viral spike protein, much remains to be understood about the molecular events behind viral dissemination. We evaluated the contribution of C-type lectin receptors (CLR_S_) of antigen-presenting cells, widely present in air mucosa and lung tissue. DC-SIGN, L-SIGN, Langerin and MGL bind to diverse glycans of the spike using multiple interaction areas. Using pseudovirus and cells derived from monocytes or T-lymphocytes, we demonstrate that while virus capture by the CLRs examined does not allow direct cell infection, DC/L-SIGN, among these receptors, promote virus transfer to permissive ACE2+ cells. A glycomimetic compound designed against DC-SIGN, enable inhibition of this process. Thus, we described a mechanism potentiating viral capture and spreading of infection. Early involvement of APCs opens new avenues for understanding and treating the imbalanced innate immune response observed in COVID-19 pathogenesis

## Introduction

C-type lectin Receptors (CLRs) are Pathogen Recognition Receptors (PRRs) involved in the detection of carbohydrate-based pathogen-associated molecular patterns by antigen-presenting cells (APC), including macrophages and dendritic cells, and in the elaboration of the immune response (Geijtenbeek and Gringhuis, 2009; Takeuchi and Akira, 2010). Many innate immune cells express a wide variety of CLRs, which differ between cell types, allowing specific adjustments of the immune response upon target recognition. Thus, CLRs such as Dectin-2, Mincle, MGL (Macrophage galactose lectin), Langerin and DC-SIGN are major players in the recognition of pathogenic fungi, bacteria, parasites and viruses (de Jong et al., 2010; van Kooyk and Geijtenbeek, 2003; Mnich et al., 2020; Van Breedam et al., 2014). The interaction of these CLRs with their ligands allows dendritic cells (DC) to modulate the immune response towards either activation or tolerance. This is done in particular through antigen presentation in lymphoid organs (primary mission of APCs) but also through the release of cytokines. Thus, DCs have a major role in modulating the immune response from the early stages of infection. To fulfill their sentinel function, DCs are localized at and patrol the sites of first contact with a pathogen, such as epithelia and mucous interfaces, including the pulmonary and nasopharyngeal mucosae. Similarly, alveolar macrophages are found in the lung alveoli.

In this battle for infection, some pathogens have evolved strategies to circumvent the initial role of CLRs in activating immunity and even to divert CLRs to their benefit for their infection process. Many viruses associate with CLRs and other host factors at the cell surface to facilitate they transfer towards their specific target receptors that will trigger fusion of viral and host membranes. This kind of viral subversion has been reported for several C-type lectin receptors, including L-SIGN (also called DC-SIGNR) and especially DC associated DC-SIGN, which promotes cis- and/or trans-infection of several viruses such as HIV, Cytomegalovirus, Dengue, Ebola and Zika viruses (Alvarez et al., 2002; Carbaugh et al., 2019; Geijtenbeek et al., 2000; Halary et al., 2002; Navarro-Sanchez et al., 2003). In particular, DC-SIGN mediates direct HIV infection of DCs (cis-infection) and can also induce trans-infection of T cells, the primary target of the virus (de Witte et al., 2008), while in the case of Dengue and Ebola, DC-SIGN allows direct cis-infection of the receptor-carrying cells (Alvarez et al., 2002; Navarro-Sanchez et al., 2003). Even more noteworthy nowadays, DC-SIGN and L-SIGN (herein after collectively referred to as DC/L-SIGN) have also been reported to be involved in the enhancement of SARS-CoV-1 infection (Jeffers et al., 2004; Marzi et al., 2004; Yang et al., 2004).

In the context of the current COVID-19 pandemic, attention is now focused on the SARS-CoV-2 virus (Huang et al., 2020; Zhou et al., 2020).Coronaviruses use a homotrimeric glycosylated spike (S) protein protruding from their viral envelope to interact with cell membranes and promote fusion upon proteolytic activation. In the case of SARS-CoV-2, a first cleavage occurs within infected cells, at the level of a furin site (S1/S2 site), generating two functional subunits S1 and S2 that remain complexed in a prefusion conformation in newly formed virus. S2 contains the fusion machinery of the virus, while the surface unit S1 contains the receptor-binding domain (RBD) and stabilizes S2 in its pre-fusion conformation. The S protein of both SARS-CoV-2 (Hoffmann et al., 2020; Letko et al., 2020; Walls et al., 2020; Zhou et al., 2020) and SARS-CoV-1 (Li et al., 2003) use ACE2 (Angiotensin-Converting Enzyme 2) as their primary receptor. For SARS-CoV-2 spike, interaction of its RBD with ACE2, as well as a second proteolytic cleavage at a S2’ site, trigger further irreversible conformational changes in S2, thus engaging the fusion process (Hoffmann et al., 2020).

The sequence of events around the S protein/ACE2 interaction are becoming increasingly clearer, but much remains to be unraveled about additional factors facilitating the infection such as SARS-CoV-2 delivery to the ACE2 receptor. Indeed, S proteins from both SARS-CoV-1 and SARS-CoV-2 have identical affinity for ACE2 (Walls et al., 2020), but this translates to very different transmission rates. We posit that the enhanced transmission rate of SARS-CoV-2 relative to SARS-CoV-1 (HCA Lung Biological Network et al., 2020) might result from a more efficient viral adhesion through host-cell attachment factors, which may promote efficient infection of ACE2^+^ cells. This type of mechanism is frequently exploited by viruses using alternatively heparan sulfate, glycolipids or CLRs to concentrate and scan cell surface for their receptor. Additionally, in the case of SARS-CoV-2, a new paradigm is needed to untangle the complex clinical picture, resulting in a vast range of possible symptoms and in a spectrum of disease severity associated on one hand with active viral replication and cell infection through interaction with ACE2 along the respiratory tract, and, on the other hand, to the development of excessive immune activation, i.e. the so called “cytokine storm”, that is related to additional tissue damage and potential fatal outcomes.

In this framework, C-type lectin PRRs and the APCs displaying them, i.e. DCs and macrophages, can play a role both as viral attachment factors and in immune activation. Thus, their role in SARS-CoV-2 infection deserves attention and we focused on DC-SIGN and L-SIGN because of their involvement in SARS-CoV-1 infections (Jeffers et al., 2004; Marzi et al., 2004; Yang et al., 2004). L-SIGN is expressed in type II alveolar cells of human lungs as well as in endothelial cells and was identified as a cellular receptor for SARS-CoV-1 S glycoprotein (Jeffers et al., 2004). DC-SIGN was also characterized as a SARS-CoV-1 S protein receptor (Marzi et al., 2004) able to enhance virus cellular entry by DC transfer to ACE2^+^ pneumocytes (Yang et al., 2004).

Recent thorough glycan and structural analyses comparing both SARS-CoV-1/2 spike glycoproteins have shown that glycosylation is mostly conserved in the two proteins, both in position and nature of the glycan exposed (Watanabe et al., 2020a, 2020b; Wrapp et al., 2020), creating a glycan shield which complicates neutralization by antibodies. Secondly, elegant molecular dynamic simulations suggested how some of the spike glycans may directly modulate the dynamics of the interaction with ACE2, stabilizing the up conformation of the RBD domain (Casalino et al., 2020; Zhao et al., 2020). Finally, and yet unexplored, spike glycans may contribute to infectivity by acting as anchor points for DC-SIGN and L-SIGN on host cells surfaces. Indeed, 28 % of glycans are of the oligomannose-type and could therefore constitutes ligands for CLRs and notably for DC-SIGN and L-SIGN. This argues also for the potential use of these CLRs by SARS-CoV-2, as do SARS-CoV-1. Additionally, some mutations modulating SARS-CoV-2 virulence have an impact on the glycosylation level of the spike. As an example, the D614G mutation, which increases virulence, has been reported as potentially increasing glycosylation at neighboring asparagine 616 (Brufsky and Lotze, 2020; Jia et al., 2020; Korber et al., 2020). A recent proteomic profiling study pointed to DC-SIGN as a mediator of genetic risk in COVID-19 (Katz et al., 2020) and finally it is of note that DC/L-SIGN expression is induced by proinflammatory cytokines such as IL-4, IL-6, IL-10 and IL-13, known to be overexpressed in severe SARS and COVID-19 cases (Lucas et al., 2020; Relloso and Puig-Kroger). These observations prompted us to investigate the potential interaction of C-type lectins receptors, notably DC/L-SIGN with SARS-CoV-2, through glycan recognition of the spike envelope glycoprotein, as well at their potential role in SARS-CoV-2 transmission.

## Results

### Production and stabilization of SARS-CoV-2 Spike Protein

In order to accurately analyze the interaction properties of the spike protein from SARS-CoV-2 with C-type lectin receptors, we expressed and purified recombinant spike protein using an expression system well-characterized in term of its site-specific glycosylation. We used here the same construct that was used 1) to obtain the cryo-electron microscopy structure of the structure (Wrapp et al., 2020) and 2) for extensive characterization of glycan distribution on the spike surface (Watanabe et al., 2020a). Expression was performed as reported, without using kifunensine to avoid blocking glycan processing. The spike protein was purified exploiting its 8xHis tag, followed by a Superose size exclusion chromatography (SEC). SEC chromatogram deconvolution allowed to select the best fractions (Figure 1B). SDS-PAGE analysis confirmed protein purity and differences in migration after reduction supported the presence of expected disulfide bridges and thus proper folding (Figure 1A). Furthermore, sample quality and trimeric assembly were confirmed by 2D class averages of the spike obtained from negatively stained sample observed under the electron microscope (Figure 1C). This construct contains “2P” stabilizing mutations at residues 986 and 987 (Pallesen et al., 2017), a inacivated furin cleavage site at the S1/S2 interface, and a C-terminal sequence optimizing trimerization (Wrapp et al., 2020). Nonetheless, we observed a limited stability over a week time scale at 4°C. To further improve protein stability and therefore ensure the quality of the following investigations, we optimized the storage buffer. Increasing ionic strength of the purification buffer up to 500 mM NaCl proved successful, preserving the trimeric state at 4°C at least for three weeks (Figure 1D to 1G). This “high-salt” concentration does not modify the structural properties of the protein as shown by identical elution profile in SEC (Figure 1B); in addition, negative-stain EM images are better in “high-salt” conditions (Figure 1D and 1 F). All protein samples were therefore subsequently produced in 500 mM NaCl and stored at −80°C. Buffer was then modified according to the analysis performed.

**Figure 1.**
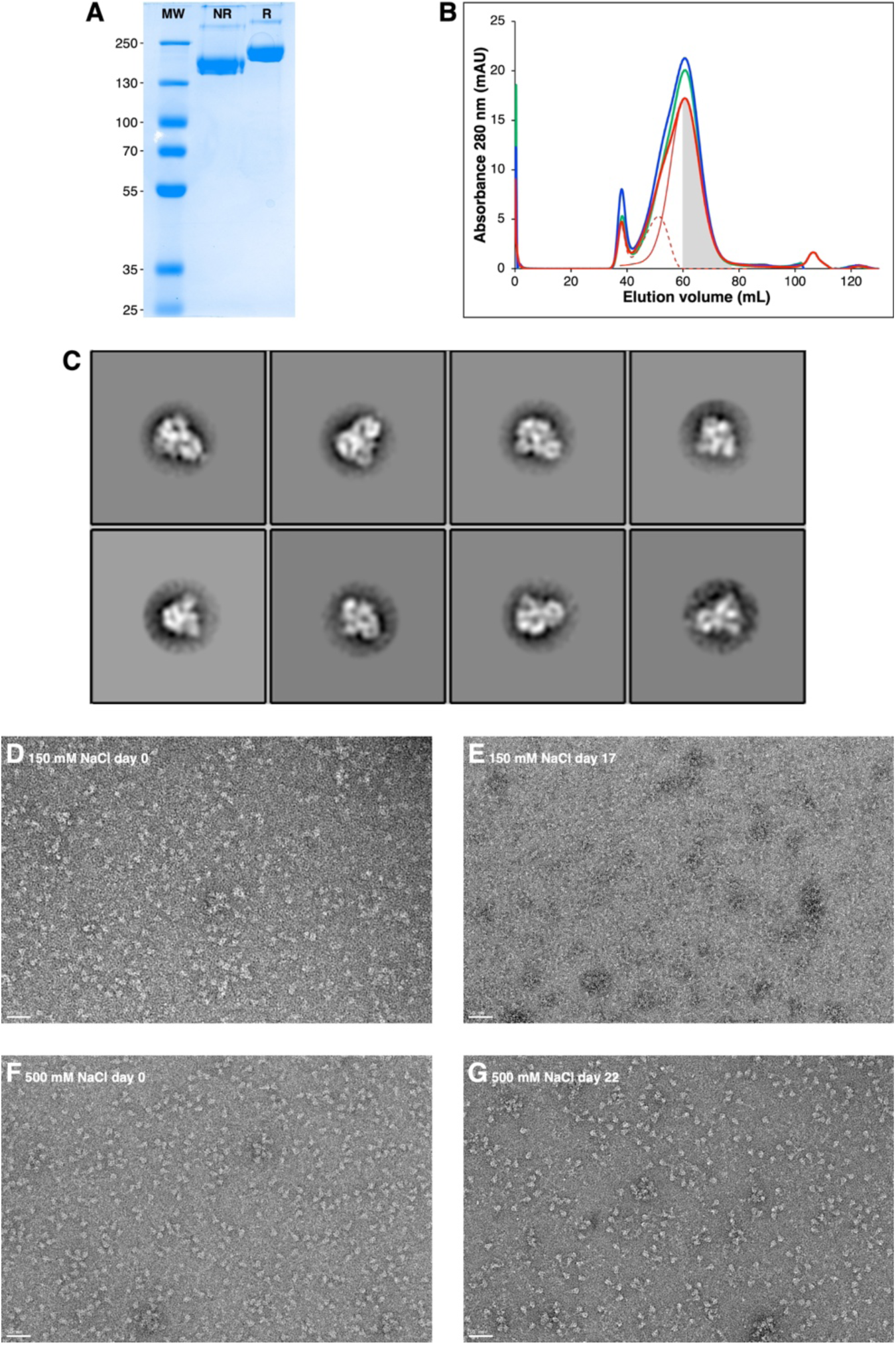
Production and optimization of spike protein. (**A**) SDS-PAGE analysis (8% acrylamide gel) of 2 µg purified SARS-Cov-2 S protein; non-reduced and reduced with mercapto-ethanol, **NR** and **R**, respectively. (**B**) Chromatograms of gel filtration profile of SARS-Cov-2 S protein using buffer with 150 mM NaCl (green line), 375 mM NaCl (blue line) and 500 mM NaCl (thick red line). Manual deconvolution of gel filtration chromatogram at 500 mM NaCl: principal peak (thin red line) and contaminants (dashed red line). Collected fractions are represented by the grey area. (**C**) Classification of 2543 particles of SARS-Cov-2 S protein after the first step of purification on HisTrap HP column, using Relion (auto-picking mode). (**D)** to (**G**) Quality control of SARS-CoV-2 S protein performed by negative staining Transmission Electron Microscopy (TEM) using uranyl acetate as stain (2% w/v). Scale bar is 50 nm. (**D)** and (**E)** Sample produced in 150 mM NaCl buffer, day of production and after 17 days at 4°C, respectively. (**F)** and (**G)** Sample produced in 500 mM NaCl buffer, day of production and after 22 days at 4°C, respectively.

### Several C-type lectin receptors can interact with SARS-CoV-2 Spike Protein

DC-SIGN and L-SIGN have been already described as receptor of SARS-CoV-1 and twenty out of the twenty-two SARS-CoV-2 S protein N-linked glycosylation sequons are conserved. Glycan shielding represent 60 to 90 % of the spike surface considering the head or the stalk of the S ectodomain, respectively (Casalino et al., 2020; Grant et al., 2020; Sikora et al., 2020). One third of N-glycans of SARS-CoV-2 spike are of the oligomannose type (Watanabe et al., 2020a). These glycans are common ligands for DC-SIGN and L-SIGN, and also for Langerin, a CLR of Langerhans cells, a subset of tissue-resident DCs of the skin, also present in mouth and vaginal mucosae (Hussain and Lehner, 1995).

To compare their recognition capabilities, SPR interaction experiments were performed with the various CLRs with immobilized SARS-CoV-2 S proteins. First, a S protein functionalized surface was generated using a standard procedure for covalent amine coupling onto the surface. The functionalization degree of this “non-oriented” surface depends upon the number of solvent exposed lysine residues (Figure 2A), which may be severely restricted by the glycan shield discussed above. Such restricted protein orientation could preclude the accessibility of some specific N-glycan clusters, located close to the linkage site and the sensor surface, thus hampering recognition by the oligomeric CLRs tested. In order to overcome these limitations, we devised and generated a so-called “oriented surface” where the S protein is captured *via* its C-terminal StrepTagII extremities onto a Streptactin functionalized surface (Figure 2B). In this set-up, no lateral parts of the S protein are attached to, and thus masked by, the sensor surface. Moreover, in the “oriented surface” set-up the spike protein is presented as it would be at the surface of the SARS-CoV-2 virus, which might better reflect the physiological interaction with host receptors. Considering both surface set-ups for the spike, the “non-oriented” one may favor access to N-glycans of the spike’s stalk domain while the “oriented” one may favor access to N-glycans of the head domain.

On the C-type lectin receptors side, we tested exclusively recombinant constructs corresponding to the extracellular domains (ECD), containing both their carbohydrate recognition domain (CRD) and their oligomerization domain. Thus, the specific topological presentation of their CRD as well as their oligomeric status is preserved for each of the CLR, going from tetramers for DC-SIGN and L-SIGN to trimers for MGL and Langerin, ensuring interactions with avidity properties as close as possible to the physiological conditions for each CLR.

Sensorgrams of interaction with both types of surface for various CLR are presented in Figure 2A and 2B. DC-SIGN and L-SIGN, initially tested on both surfaces, recognized the spike with the same profile, whatever the set-up. Thus, the next two CLRs were tested only on one type of surface. Langerin was found to interact with the S protein in agreement with the presence of oligomannose-type glycans. Finally, MGL, a lectin that specifically recognizes glycans bearing terminal Gal or GalNAc residues, also interacted with the S protein (Figure 2A). This shows that complex N-glycans may also serve as potential anchors for the SARS-CoV-2 S protein to cell surface CLRs.

**Figure 2.**
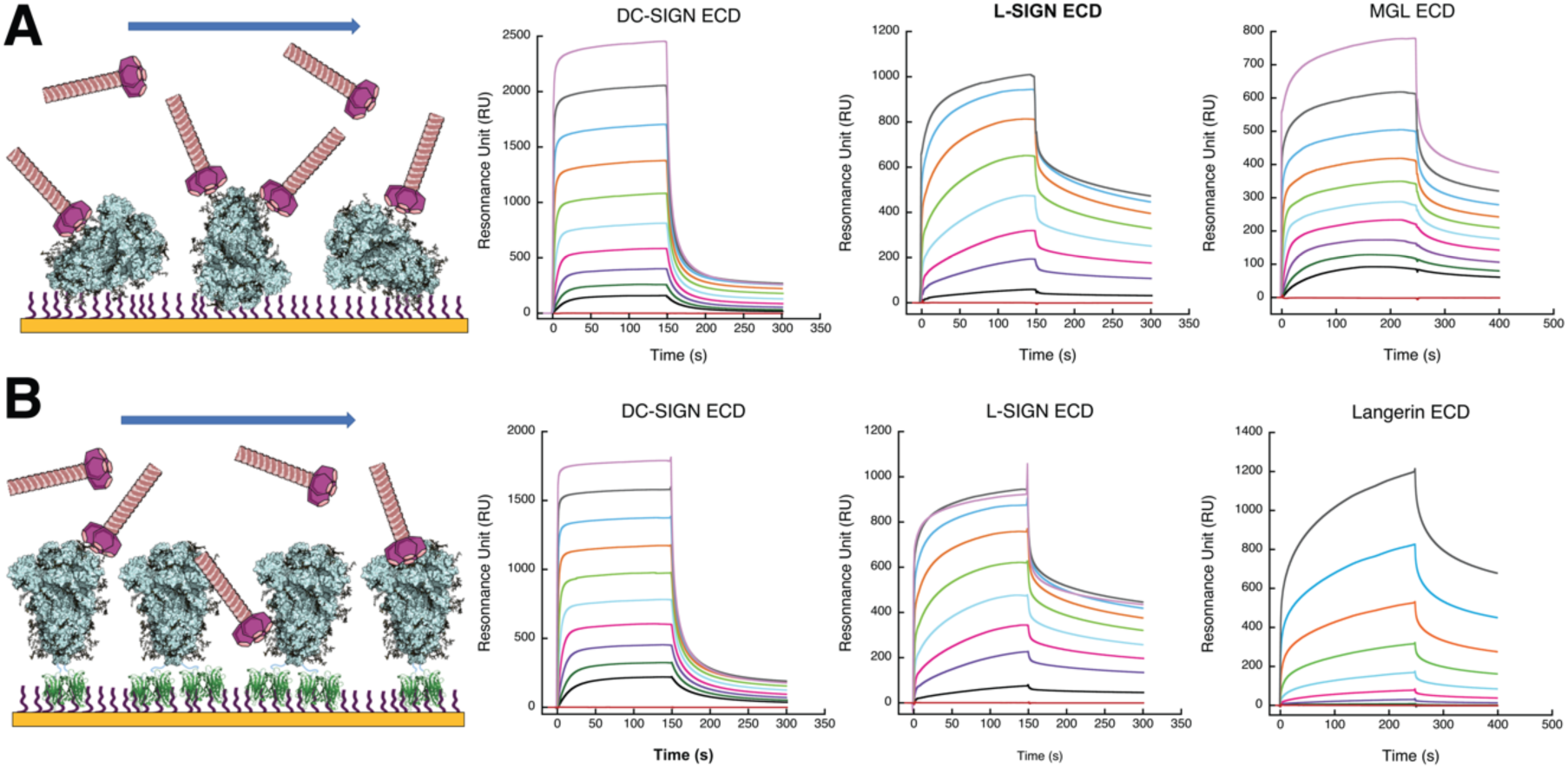
C-type lectin interactions with SARS-CoV-2 S protein. **(A)** Covalently functionalized S protein surface in random orientation. Multivalent C-type lectins oligomeric ectodomains (ECD) are injected over the surface. From left to right, surfaces titrations were performed with DC-SIGN ECD (tetrameric), L-SIGN ECD (tetrameric) and MGL-ECD (trimeric). (**B)** Surfaces were first functionalized with streptactin (in green) and then S protein was captured by its C-terminal double-StrepTagII. From left to right, surfaces titrations were performed with DC-SIGN ECD (tetrameric), L-SIGN ECD (tetrameric) and Langerin-ECD (trimeric). The same color code for concentrations of all CLR-ECD has been used for all panels. Concentrations injected (i.e DC-SIGN ECD in A) range from 50 µM to 0.1 µM by 2-fold serial dilutions (decreasing concentrations from top to bottom). The red sensorgram, lower flat line, correspond to control experiments with buffer injection deprived of CLR.

While all CLRs tested interacted with the spike, the interactions observed are not all equivalent. Unfortunately, the complexity of the process involving probably multiple binding sites per oligomeric CLR prevented a kinetic fitting using classical kinetic models, which precluded the determination of kinetic rate constants. Nevertheless, an apparent equilibrium dissociation constant (*K*_D_) could be obtained by steady state fitting for DC-SIGN, L-SIGN and MGL. For Langerin, despite a longer injection time, a much higher range of concentration would have been required to reach the equilibrium and accurately evaluate an apparent *K*_D_. DC/L-SIGN and MGL showed affinities in the µM range, from around 2 to 10 µM (Table 1), depending on the CLR and the surface type, while Langerin has an affinity of at least one order of magnitude lower. Despite the impossibility to evaluate kinetic association and dissociation rate constants (*k*_*on*_ and *k*_*off*_), visual inspection of the sensorgrams clearly reveals different behaviors between DC-SIGN and L-SIGN independent of the surface set-up. While association and dissociation seem to be very fast for DC-SIGN, L-SIGN sensorgrams suggest a much slower association and dissociation rate that compensate each other to provide a *K*_D_ similar to that of DC-SIGN. However, while the higher *k*_on_ value for DC-SIGN argues for a faster formation of the DC-SIGN/S protein complex, the lower *k*_*off*_ value for L-SIGN suggests that the L-SIGN/S-protein complex might be more stable. Finally, for DC-SIGN and L-SIGN, which have been tested both on “non-oriented” and “oriented” S surface, no obvious differences has been observed in the interaction sensorgrams. This suggests that the interaction is not restricted to a limited glycan cluster, but rather that oligomannose-type glycans are multiple, accessible and distributed over the whole S protein.

**Table 1:** Steady state determination of apparent *K*_D_ for CLRs interaction with SARS-CoV-2 S protein.

|  | DC-SIGN | L-SIGN | MGL | Lang |
| --- | --- | --- | --- | --- |
| | $\mu\text{M}$ | $\mu\text{M}$ | $\mu\text{M}$ | |
| Non-oriented | $11.90 \pm 4.6$ | $1.80 \pm 0.12$ | $7 \pm 1.5$ | n.a. |
| Oriented | $4.33 \pm 0.7$ | $1.66 \pm 0.12$ | - | - |

### DC-SIGN forms multiple complexes with SARS-CoV-2 Spike Protein

The SPR interaction analysis argues for multiple potential binding sites for CLRs on the S protein. Such initial host adhesion mechanism could be essential for efficient viral capture, viral particles concentration on the cell surface and subsequent enhanced ACE2 targeting and infection. Negative stain electron microscopy was used to visualize potential DC-SIGN/S protein complexes.

Extemporaneously after a fresh purification of S protein, SEC fractions corresponding to the pure trimeric spike were mixed with a DC-SIGN ECD preparation in a molecular ratio 1:3 (meaning 1 trimeric spike for 3 tetrameric DC-SIGN ECD). In order to enrich the proportion of complex in the sample for EM observation, we directly reinjected this mix onto same SEC column and recovered fractions in the elution profile corresponding to higher molecular weight, thus potentially corresponding to DC-SIGN/S protein complex. These fractions were immediately used to prepare and observe negatively stained electron microscopy grid (Figure 3). Figure 3A and 3B show control images of the DC-SIGN and spike sample used and images of the sample corresponding to the enriched complexes are shown in Figure 3C where DC-SIGN/S complexes can be clearly seen. Most images show a 1:1 complex but, in some cases, (Figure 3C, left) at least two molecules of DC-SIGN interacting with the spike protein can be detected. Even if it is not possible to precisely define the interacting region on the spike protein, several areas seem to be involved in the interaction with DC-SIGN.

**Figure 3.**
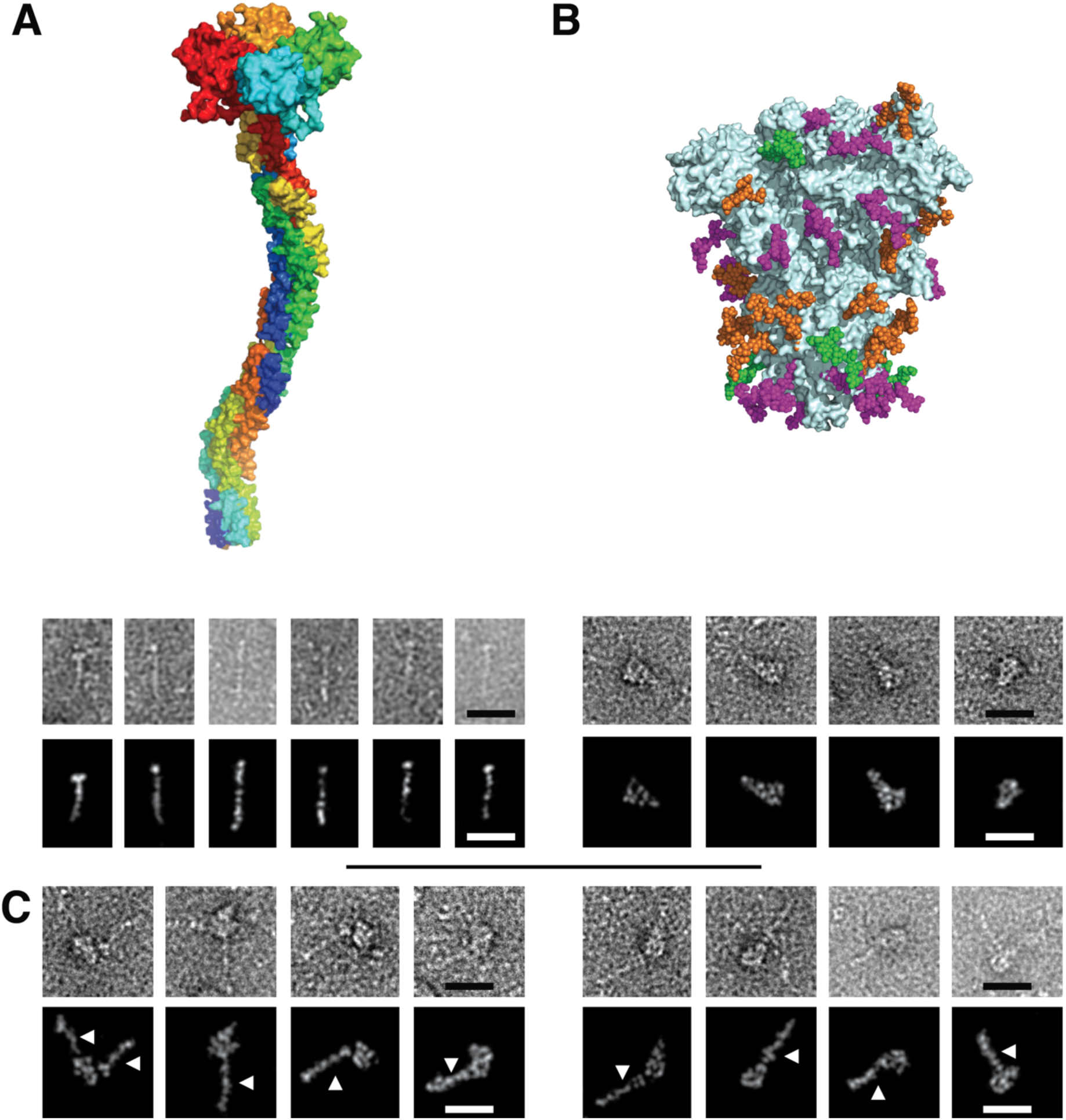
Electron Microscopy micrographs of DC-SIGN/S protein complexes. **(A)** DC-SIGN. Top: model of DC-SIGN ECD tetramer adapted from Tabarani et al (2009). On the bottom: Negative staining images of DC-SIGN. Top row: original images; bottom row: Photoshop processed images. The scale bar represents 25 nm. (**B)** Spike protein. Top: model of the glycosylated Spike adapted from model of Casalino et al pdb 6vsb). Glycan sites are represented with color code derived from the work of Crispin *et coll*. (Watanabe et al., 2020a), according to oligomannose-type glycan content, in green (80-100%), orange (30-79%) and magenta (0-29%). On the bottom: Negative staining images of spike protein. Top row: original images, bottom row: Photoshop processed images. The scale bar represents 25 nm. **(C)** Complex between DC-SIGN and spike protein. Negative staining image of the complexes between DC-SIGN and spike protein. The white arrows highlight DC-SIGN molecules. Top row: original images; bottom row: Photoshop processed images. The scale bar represents 25 nm.

### Antigen-Presenting Cells expressing DC-SIGN, MDDCs and M2-MDM are not infected by SARS-CoV-2 pseudovirions

To study the potential role of DC/L-SIGN in SARS-CoV-2 infection, VSV/SARS-CoV-2 pseudotyped viruses were firstly used for direct infection of primary monocyte-derived cells, including monocyte-derived DCs (MDDCs) and M2 monocyte-derived macrophages (M2-MDM), that have been shown to express DC-SIGN. As a control, VSV/EBOV-GP pseudotypes, expected to display enhanced infection in the presence of DC/L-SIGN receptor was used as well as VSV/VSV-G as a DC/L-SIGN independent pseudotype. To evaluate the infection mediated exclusively by DC/L-SIGN receptor on primary monocyte-derived cells we also examined the infection with these three pseudotypes in the presence of anti-DC/L-SIGN antibody.

VSV/SARS-CoV-2 did not infect MDDCs or M2-MDMs, despite DC-SIGN expression (Figure 4A). As expected, all primary cell lines were efficiently infected with VSV/EBOV-GP and this infection could be substantially blocked by an antibody targeting DC-SIGN. Infection of primary cells with EBOV does not exclusively depend on the presence of DC-SIGN on the cell surface, since other receptors are also responsible for the direct infection with EBOV (Marzi et al., 2007) which explains the residual infection observed in the presence of an anti-DC/L-SIGN. DC-SIGN-mediated cis-infection was clearly blocked with anti-DC/L-SIGN in the case of MDDCs (92.5% inhibition of infection), followed by M2-MDM (68.4% inhibition). VSV/VSV-G also showed a great infectivity of all primary cell lines. However, this infection was DC-SIGN independent, since anti-DC-SIGN antibodies did not impact the infection level (Figure 4A).

**Figure 4.**
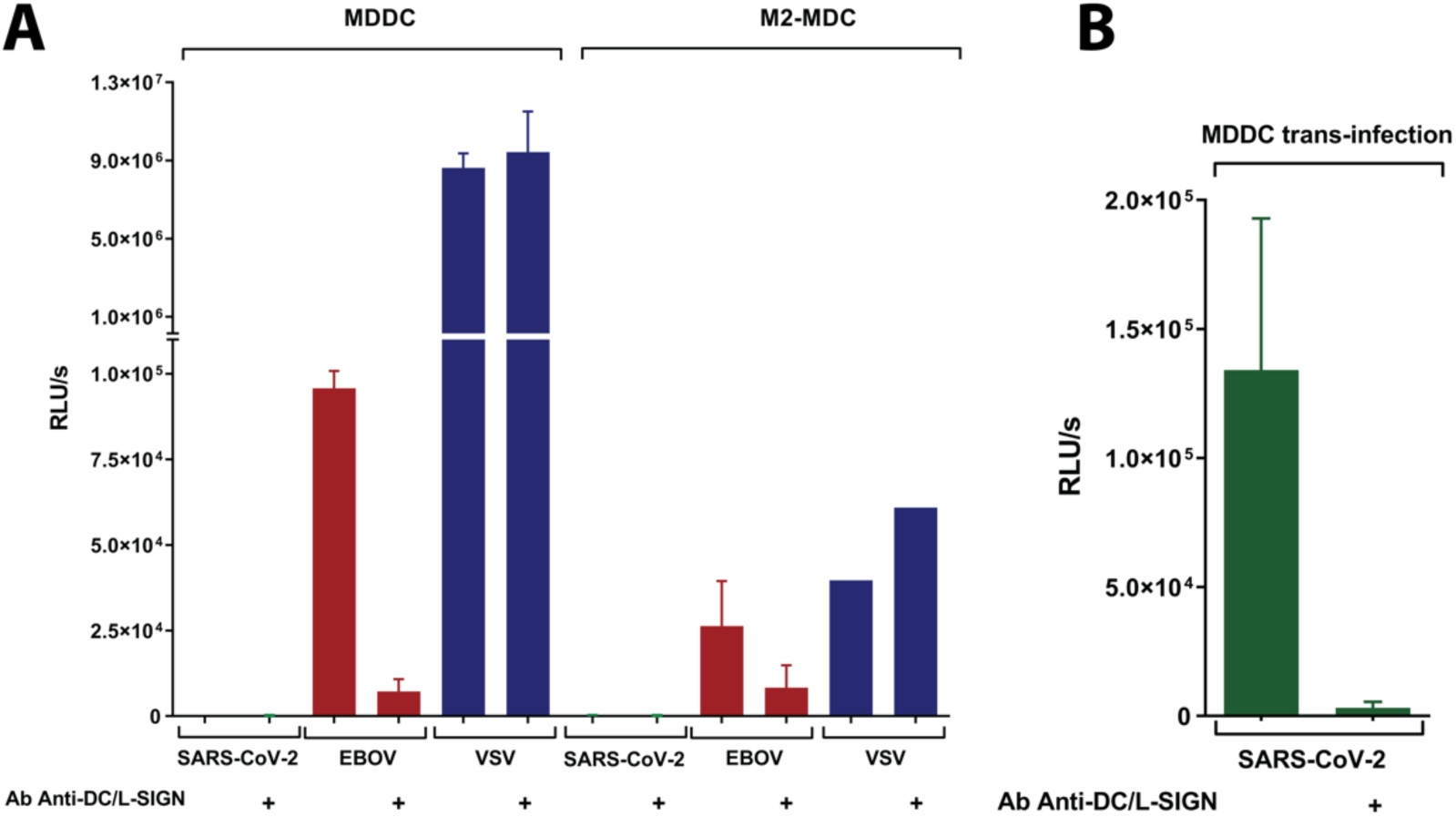
SARS-CoV-2 *cis-* and *trans*-infection of monocytes-derived antigen presenting cells. (**A)** Direct infection of primary cells: monocyte-derived dendritic cells (MDDCs), and M2 monocyte-derived macrophages (M2-MDM). Cells (5 × 10^4^) were challenged with SARS-CoV-2, EBOV-GP or VSV-G pseudotyped recombinant viruses. Bars represent mean ± SEM of mean of three independent experiments with cells from different donors performed in duplicates. As a control of inhibition of infection mediated by DC-SIGN, anti-DC-SIGN Ab was used. (**B**) Trans-infection using MDDCs. Cells (5 × 10^4^) were challenged with SARS-CoV-2 pseudotyped recombinant viruses during 2 h at room temperature with rotation and then were co-cultivated with adherent Vero E6 cells for 24 h. Bars represent mean ± SEM of mean of three independent experiments with cells from different donors performed in duplicates. As a control of inhibition of trans-infection mediated by DC-SIGN, anti-DC-SIGN Ab was used. Infection values from luciferase assays are expressed as Relative Light Units (RLUs).

### MDDCs promote DC-SIGN-mediated trans-infection of SARS-CoV-2 pseudovirions

DC/L-SIGN are known to enhance viral uptake for direct infection in the process referred to as cis-infection and can also internalize viral particles into cells for storage in non-lysosomal compartments and subsequent transfer to susceptible cells in the process recognized as trans-infection (Alvarez et al., 2002; Geijtenbeek et al., 2000). To study the potential function of DC/L-SIGN in SARS-CoV-2 trans-infection, MDDCs were incubated with VSV/SARS-CoV-2 for 2 h and, after extensive washing, they were placed onto susceptible Vero E6 cells, the reference ACE2+ cell line for SARS-CoV-2 cell culture (Zhou et al., 2020). Interestingly, DC-SIGN promoted efficient SARS-CoV-2 trans-infection from MDDC to Vero E6 (Figure 4B). An Anti-DC-SIGN antibody could reduce substantially the infectivity observed (98% inhibition), confirming the role of this CLR in the process of SARS-CoV-2 trans-infection.

### DC/L-SIGN but not Langerin mediate trans-infection of SARS-CoV-2 in a T-lymphocyte cell line

The role of DC/L-SIGN and Langerin in SARS-CoV-2 infection was also examined in Jurkat cells (a T-lymphocyte cell line lacking ACE2 expression). Parental Jurkat cell line, which does not express DC-SIGN, was used as control of infectivity along with Jurkats stably expressing DC/L-SIGN and Langerin (Alvarez et al., 2002). VSV/VSV-G pseudovirions were used in parallel. VSV-G cell entry is independent of the presence of CLRs and can efficiently infect Jurkat cells. Direct infection assay with VSV/SARS-CoV-2 showed no infectivity of any of the cell line tested, Jurkat, Jurkat DC-SIGN and Jurkat L-SIGN (Figure 5A), indicating that DC/L-SIGN do not function as direct receptors for SARS-CoV-2. As expected, VSV/VSV-G could infect all of the cell lines at similar level. The use of DC/L-SIGN and Langerin by SARS-CoV-2 was further verified in trans-infection assays using Jurkat, Jurkat DC-SIGN, Jurkat L-SIGN and Jurkat Langerin cells. As controls, VSV/EBOV-GP and VSV/VSV-G were used in parallel. VSV/EBOV-GP trans-infection based on Jurkat cells is absolutely dependent on the presence of DC/L-SIGN or Langerin on the cell surface, since EBOV does not infect T-lymphocytes (Yang, 1998). On the other hand, VSV/VSV-G is DC/L-SIGN or Langerin independent. Jurkat, Jurkat DC-SIGN, Jurkat L-SIGN and Jurkat Langerin cells were incubated with VSV-based pseudotypes for 2h and after extensive washing, they were co-incubated upon susceptible Vero E6 cells monolayers. Similarly to the results obtained with primary cells, we could observe that DC/L-SIGN, but not Langerin, promoted efficient SARS-CoV-2 trans-infection of Vero E6 (Figure 5B). Trans-infection in the presence of antibodies anti-DC/L-SIGN could significantly reduce DC/L-SIGN-mediated trans-infection (86.7% and 78.7% inhibition of transinfection, respectively) confirming the important role of these receptors in the SARS-CoV-2 trans-infection process.

Interestingly, no trans-infection was observed using Jurkat Langerin cells (Figure 5B), which could indicate that Langerin does not serve as SARS-CoV-2 trans-infection receptor. As expected, VSV/EBOV-GP achieved high DC/L-SIGN-mediated trans-infection of Vero E6 cells which could be blocked by anti-DC/L-SIGN antibody (96.9% and 97.3% inhibition of infection respectively). VSV/EBOV-G could also use Langerin as an attachment factor for trans-infection and this process could be partially blocked by anti-Langerin antibody (57.2% inhibition of infection). On the other hand, VSV/VSV-G does not use CLRs for trans-infection, thus no transmitted infection was detected in Vero E6 cells. The ratio of trans-infection mediated by DC/L-SIGN was calculated for VSV/SARS-CoV-2, VSV/EBOV-GP and VSV/VSV-G using the parental Jurkat cell line as reference. The ratio of VSV/SARS-CoV-2 trans-infection using Jurkat DC-SIGN and Jurkat L-SIGN was 62% and 35% respectively. VSV/EBOV-GP trans-infection ratio was 23% for Jurkat DC-SIGN and 172% for Jurkat L-SIGN. VSV/VSV-G did not show any appreciable DC/L-SIGN mediated trans-infection (Figure 5C).

**Figure 5.**
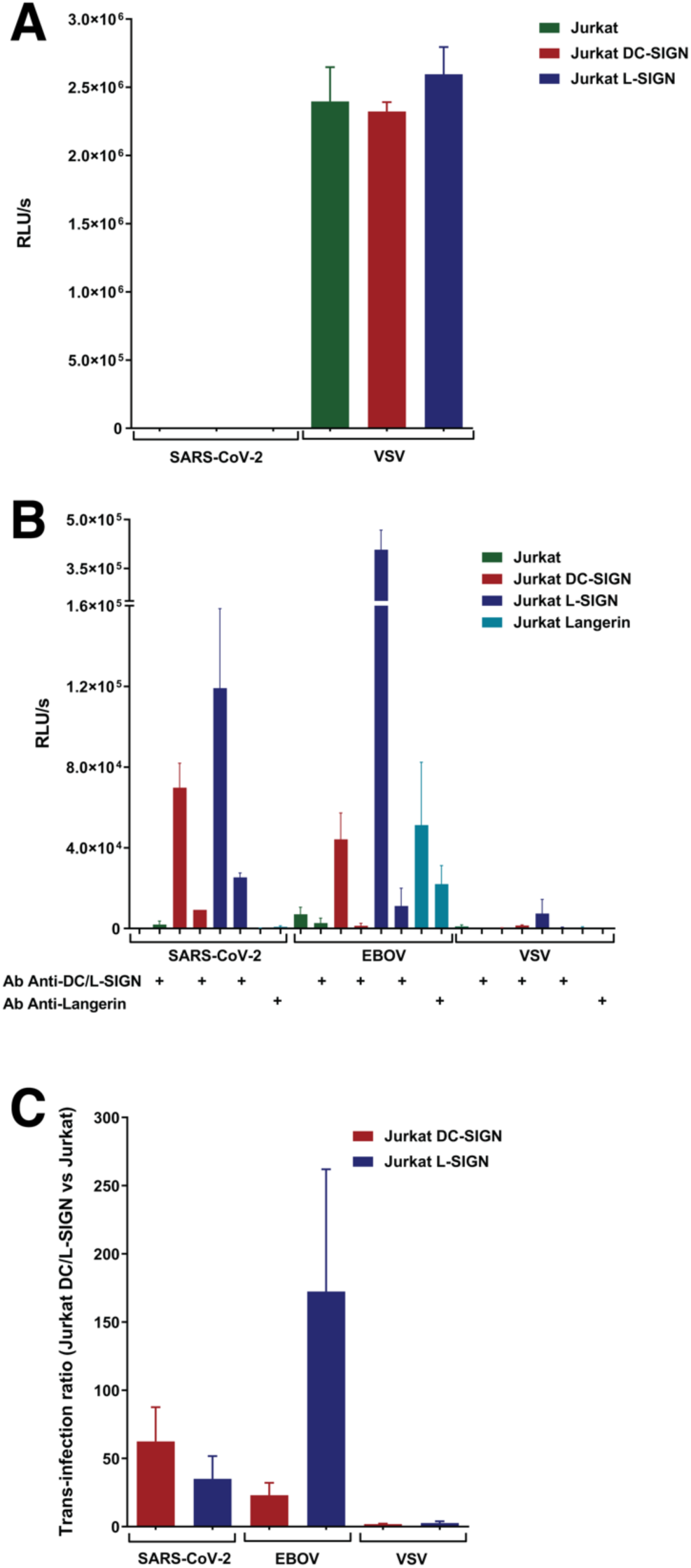
Impact of DC/L SIGN and Langerin in cis- and trans-infection of SARS-CoV-2. (**A)** Direct infection of Jurkat, Jurkat DC-SIGN and Jurkat L-SIGN with SARS-CoV-2 and VSV-G pseudotyped recombinant viruses. Cells (5 × 10^4^) were challenged with SARS-CoV-2, or VSV-G pseudotyped recombinant viruses. (**B)** Trans-infection using Jurkat cell with or without CLR DC-SIGN, L-SIGN and Langerin. As a control of inhibition of trans-infection mediated by CLRs, Ab specifically directed toward the corresponding CLR were used. (**C)** Trans-infection ratio of Jurkat DC-SIGN or Jurkat L-SIGN versus Jurkat cells. Cells were challenged with SARS-CoV-2, EBOV and VSV-G pseudotyped recombinant viruses during 2 h at room temperature with rotation and then were co-cultivated with adherent Vero E6 cells for 24 h. Bars represent mean ± SEM of mean of two independent experiments performed in triplicates. Infection values from luciferase assays are expressed as Relative Light Units (RLUs) and ratios represents RLUs obtained by trans-infection with Jurkats expressing CLRs divided by those obtained from parental Jurkats.

### DC-SIGN binding to the S protein and DC-SIGN-dependent *trans*-infection are inhibited by a known glycomimetic ligand of DC-SIGN (PM26)

Polyman26 (PM26, Figure 6A) is a multivalent glycomimetic mannoside tailored for optimal interaction with DC-SIGN (Ordanini et al., 2015). It is known to bind DC-SIGN carbohydrate recognition domain (CRD), eliciting a Th-1 type response from human immature monocyte derived dendritic cells (Berzi et al., 2016). It also inhibits DC-SIGN mediated HIV infection of CD8^+^ T lymphocytes with an IC_50_ of 24 nM (Ordanini et al., 2015).

PM26 was used in SPR competition experiments to inhibit DC-SIGN binding to immobilized spike protein, both in the oriented and non-oriented set-ups (Figure 6B-C). The lectin (20 µM) was co-injected with variable concentrations of PM26 (from 50 µM to 0.1 µM), and the results showed clear dose-dependent inhibition. No significant difference was observed between the oriented and non-oriented surface, which is consistent with the binding data previously discussed (Figure 2). Thus, an IC_50_ of 9,6±0,4 µM correlates with the interaction affinity between DC-SIGN and the spike functionalized surfaces. It suggests, in such competition test were the reporting interaction can be limiting (Porkolab et al., 2020), that a real higher avidity towards DC-SIGN can be awaited for PM26 (Figure 6C).

**Figure 6.**
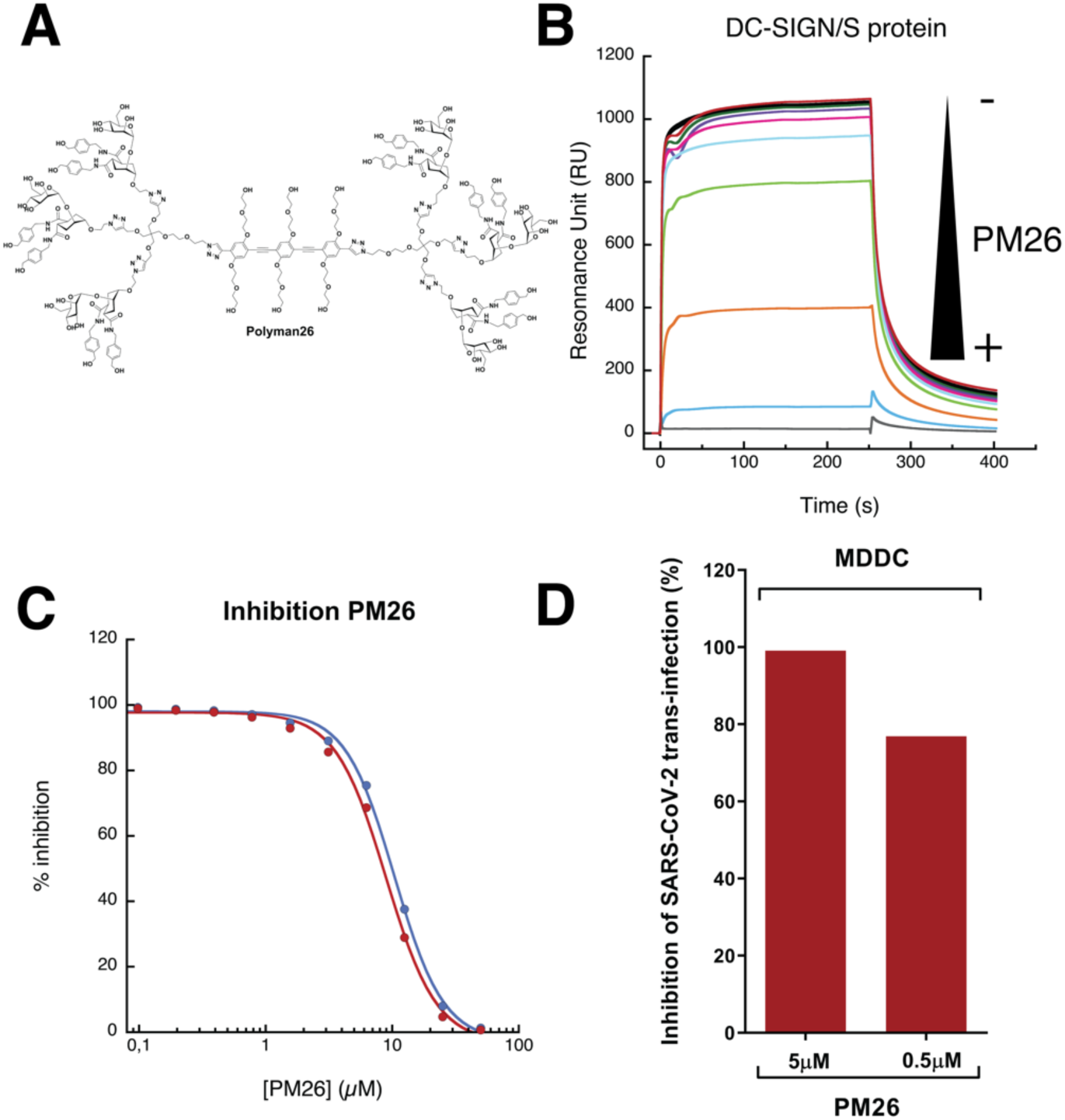
PM26, a specific antagonist of DC-SIGN, can inhibit DC-SIGN/S protein interaction and trans-infection of SARS-CoV-2. **(A)** Structure of Polyman26 (PM26). **(B)** Sensorgrams of DC-SIGN/S protein interaction inhibition by a range of PM26 concentration. DC-SIGN is injected at the constant concentration of 20 µM, PM26 is co-injected at decreasing concentration going from 50 µM to 0.1 µM. S protein is presented in an oriented mode in the experiment presented here. **(C)** PM26 inhibition curve of the DC-SIGN S protein interaction using an oriented S protein surface (blue) or non-oriented S protein surface (red) set up as described in Figure 2. **(D)** Inhibition of trans-infection of MDDCs with SARS-CoV-2. MDDCs were incubated for 20 min with PM26 at 5 and 0.5 μM before being challenged with SARS-CoV-2 for 2h at rotation. After extensive washing, MDDCs were co-cultivated with susceptible Vero E6 for 24h. Bars correspond to a single experiment of inhibition of SARS-CoV-2 trans-infection. Results are presented as percentage of inhibition of SARS-CoV-2 trans-infection in the presence of PM26 as compared to MDDCs mediated SARS-CoV-2 trans-infection of susceptible Vero E6 without addition of any compound.

The possibility of blocking DC-SIGN-mediated SARS-CoV-2 trans-infection using PM26 was also examined. DCs were pre-incubated with PM26 for 20 min before being challenged with VSV/SARS-CoV-2. The results of blocking DC-SIGN receptor by PM26 are shown as a percentage of inhibition of infection transmitted by DCs to susceptible Vero E6, as compared with SARS-CoV-2 trans-infection mediated by DCs to Vero E6 without addition of any compound. PM26 was tested at two different concentrations: 5 and 0.5 μM showing 99% and 77% of inhibition of SARS-CoV-2 trans-infection, respectively, which is consistent with the results described for HIV inhibition and confirming an effective affinity in the nanomolar range for PM26 (Ordanini et al., 2015).

## Discussion

Even if viruses target mainly one specific cellular entry receptor within their infection cycle, their efficiency often largely depends upon additional binding events at the cell surface, which promote access to the so-called primary receptor. Although such additional receptors may not promote any fusion step, they can drive viral internalization through endocytic processes or simply by viral adhesion to the host cell, accumulation of viral particles on the cell surface and finally engagement with the primary receptor followed by the fusion event. Different types of attachment factors can be found on the host cell surface: either glycans, such as heparan-sulfate, glycolipids or protein N-glycans, often targeted by envelope viral protein with lectin-like properties (Dimitrov, 2004), or immune lectin-type receptors including CLRs and Siglecs (Bermejo-Jambrina et al., 2018; Perez-Zsolt et al., 2019). CLRs primary role is to sense the presence of pathogen through recognition of specific non-self glycans. However, some viruses are able to hijack them as co-receptors for cell entry or hiding. HIV represents a well-known example since it exploits DC-SIGN in genital mucosa to promote uptake by DCs and T lymphocytes trans-infection (Geijtenbeek et al., 2000).

SARS-CoV-1 and MERS-CoV have been described to use heparan sulfate and sialic acid exposed at the cell surface to attach to cell membranes (Milewska et al., 2014; Park et al., 2019; Tortorici et al., 2019). Remarkably, DC/L-SIGN were not only reported to act as attachment factors for SARS-CoV-1, but also as promoters of viral dissemination through trans-infection. Here we investigate whether those CLRs could also play a role in SARS-CoV-2 infection and dissemination. While we were finalizing this work, a first preprint article described the binding (Elisa test) of Fc-CLR constructs, notably based on DC-SIGN and MGL, to a commercial RBD domain of the SARS-CoV-2 (Chiodo et al., 2020). Given the importance of the role played by glycan determinants in this recognition event, therefore peculiar attention must be paid to the quality of the S protein sample used. Indeed, it has to be ideally as close as possible to the physiological product in terms of glycosylation pattern and distribution. In particular, the expression system considered as well as the local protein environment may have a strong impact on the type of glycan added as well as on their level of maturation. Viral envelope proteins display a dense array of glycans resulting from evolutionary pressure to mask immunogenic epitopes at their surfaces. This glycan density coupled to specific structural features of envelope proteins generate steric constraints preventing proper access of glycan processing enzymes to some substrate glycans (Behrens and Crispin, 2017). Expressing the whole spike ectodomain or just the single RBD domain may therefore lead to very different N-glycan distribution, especially considering that the RBD contains only 2 N-glycosylations sites, while up to 66 N-glycan sites are found over the whole spike protein. For these reasons, we selected the entire ectodomain of S as our model to investigate additional attachment factors for SARS-CoV-2. We expressed the protein using the same construct enabling the spike EM structure (Wrapp et al., 2020) and its glycan profiling (Watanabe et al., 2020a), using a HEK293-derived expression system known to provide glycosylation pattern similar to epithelial tissues. Similarly, we expressed the entire ectodomain for the CLRs as well, avoiding Fc-CRD constructs, in order to preserve their specific oligomeric assembly and therefore their avidity properties.

Using SPR we showed that all the C-type lectin receptors tested interact with the spike protein. Three of those, DC-SIGN, L-SIGN and langerin share the ability to recognize high-mannose oligosaccharides. In particular, L-SIGN is tightly specific for high-mannose, while DC-SIGN additionally recognizes fucose based ligands (several Lewis type glycans) and Langerin binds sulfated sugars as well. MGL is specific for Gal and GalNAc terminated glycans and may bind to complex N-glycans as a function of their level of maturation (Valverde et al., 2020). Analyzing the glycosylation pattern of the spike protein, reported in Figure 3B, all glycosylation site depicted in green or orange are potential ligands for L-SIGN and Langerin, with different level of probability from site to site, while MGL’s ligands will be found in magenta sites. DC-SIGN might potentially recognize all of them. Beside all considerations about specificity, the accessibility of N-glycan sites upon spike presentation at the SARS-CoV-2 virus surface is also of paramount importance for recognition. DC-SIGN and L-SIGN share the same tetrameric organization and they recognize with similar avidity the spike functionalized SPR surface, suggesting that they share a primary recognition epitope - i.e. high mannose. The SPR experiments described here have been performed sequentially on the same spike surfaces with the different CLRs. Of these, DC-SIGN and L-SIGN have similar organization and molecular weight (Feinberg et al., 2001), thus the difference in RU level reached by the two lectins at their equilibrium (approx. 1000 RU higher for DC-SIGN) suggests that there is more DC-SIGN binding and thus more epitopes available for it, implying that high mannose are not the unique glycan epitope used here by DC-SIGN. This suggest that DC-SIGN may be able to also bind to some o complex N-glycosylation sites (in magenta in Figure 3B), possibly presenting a proper fucosylation pattern that generates Lewis-type epitopes. These considerations, in addition to the oligomeric state of the CLRs examined, lead us to rule out a simple interaction with a single preferential epitope and a 1:1 stoichiometry in favor of a more complex picture with multiple and simultaneous binding events, much like the “Velcro effect” often recalled when discussing glycan-protein interactions. This is clearly supported by the EM characterization of Spike/DC-SIGN complexes (Figure 3C) that shows several interactions areas on the spike and can also explain the absence of affinity differences between non-oriented and oriented spikes surfaces in SPR.

The complexity of the binding event(s) described above does not allow to extract kinetic association and dissociation rates from the sensorgrams. Only a global apparent *K*_D_ could be inferred, giving avidity levels. However, L-SIGN may have a slightly better affinity (around 2 µM, while values ranging from 5 to 10 µM have been obtained for DC-SIGN and MGL) and seems to generate more stable complexes. Such µM range of affinity, as determined here for soluble forms of CLR, will result in surface avidity of several order of magnitude higher at the cell membrane (Porkolab et al., 2020). Indeed, CLRs, as DC SIGN, are organized in microdomain allowing multiple attachment point for viral capture (Cambi et al., 2004).

CLRs and particularly DC/L-SIGN have been associated to important steps of viral entry and infection of different viruses. Participation of DC-SIGN in the infectivity and initial dissemination of a number of viral agents has been described in animal models for measles (Mesman et al., 2012, 2014), Japanese encephalitis virus (Liang et al., 2018) and *in vivo* for HIV-1. The founder viruses that initiate HIV infection through mucosa exhibit higher content of high-mannose carbohydrates (Go et al., 2011), as well as higher binding to DCs dependent on DC-SIGN expression (Parrish et al., 2013). In the case of Ebola virus, DCs and macrophages have been shown to be the initial targets of infection in macaques (Geisbert et al., 2003; Martinez et al., 2012) and circulating DC-SIGN^+^ DCs have been shown to be the first cell subset infected upon intranasal EBOV inoculation in a murine model (Lüdtke et al., 2017).

In SARS-CoV-1 infection it was shown that DC/L-SIGN can enhance viral infection and dissemination (Marzi et al., 2004; Yang et al., 2004) and even it has been proposed that L-SIGN could act as an alternative cell receptor to ACE2 (Jeffers et al., 2004). Our work shows that DC/L-SIGN are important enhancers of infection mediated by the S protein of SARS-CoV-2 that greatly facilitate viral transmission to susceptible cells. *In vivo*, DC-SIGN is largely expressed in immature dendritic cells in submucosa and tissue resident macrophages, including alveolar macrophages (Tailleux et al., 2005) whereas L-SIGN is highly expressed in human type II alveolar cells and lung endothelial cells (Jeffers et al., 2004). Using primary MDDCs and M2-MDM, two of the established primary cell models to explore DC-SIGN interactions (Dominguez-Soto et al., 2007) and a well-established VSV-based pseudovirion system, we did not observe direct infection of MDDCs or M2-MDM, indicating that DC-SIGN expressed by these cells does not function as an alternative receptor (Figure 4A). MDDCs and M2-MDM, as expected, were infected by VSV pseudotyped with EBOV-GP or VSV-G, although in a DC-SIGN dependent and independent manner respectively. DC-SIGN, however, showed a clear function as an enhancer of infection, in a process known as trans-infection (Geijtenbeek et al., 2000). An obvious increase of infection was observed when SARS-CoV-2 pseudovirions were incubated with these primary cells and then placed in contact with susceptible VeroE6 cells (Figure 4B). Similar results were obtained with the T lymphocyte Jurkat cell line. T lymphocytes lack ACE2 expression (Hamming et al., 2004) and both the parental Jurkat cell line and Jurkats expressing DC/L-SIGN were not directly infected by SARS-CoV-2 pseudovirions (Figure 5A). Therefore, we did not observe that these CLRs can function as alternative receptors to ACE2 in non-permissive cells such as T lymphocytes or HEK 293 (Supplementary information), as it has been recently suggested by Amraie et al. (Amraie et al., 2020). On the other hand, DC/L-SIGN expression on Jurkat cells allows binding and capture of SARS-CoV-2 pseudovirions. This trans-infection mechanism is significantly inhibited by a specific DC/L-SIGN antibody and, remarkably, it seems exclusive of DC/L-SIGN, since the related CLR langerin does not mediate trans-infection in Jurkat cells. This is similar to what has been reported for HIV-1, since langerin acts as an antiviral receptor that degrades HIV-1 via internalization and subsequent degradation (de Witte et al., 2007). On the other hand, langerin appears to function as a trans-receptor for EBOV pseudovirions highlighting the complexity of CRLs recognition patterns and functionalities.

The biological relevance of the interaction of CLRs with SARS-CoV-2 remains yet to be fully explored. The innate immune system provides the first line of defense against viruses and it is now clear that severe COVID-19 is largely due to an imbalance between viral replication and antiviral and pro-inflammatory responses (Yang et al., 2020). Here we demonstrate that DC/L-SIGN can function, in primary cells and cell lines, as potent trans-receptors. The expression of DC/L-SIGN in relevant cell subsets along the respiratory tract, such as submucosal DCs and Macrophages or type II alveolar cells together with their potency to enhance viral infection could be critical for the pathogenesis of COVID-19. In this context, it is important to note that DC-SIGN expression is negatively regulated by IFN, TGF-β, and anti-inflammatory agents (Relloso et al., 2002) and that DC/L-SIGN activation though ligand recognition triggers an immune response consisting in the production of pro-inflammatory cytokines such as IL-6 and IL-12 (Bermejo-Jambrina et al., 2018). These cytokines are raised up in severe forms of COVID-19 and are among the mediators responsible for the development of the cytokine storm and the hyper-inflammatory syndrome related to poor prognosis and eventual fatalities(Vabret et al., 2020). DC/L-SIGN expression can be upregulated as well, since it has been demonstrated that while innate immune responses are potently activated by SARS-CoV-2, it also counteracts the production of type I and type III interferon (Blanco-Melo et al., 2020).

Innate immunity plays an important role in initial control and the pathogenesis of COVID-19. Indeed, there are a number of clinical trials focused on the use of interferon and anti-inflammatory agents to counteract the sometimes deleterious immune response to SARS-CoV-2 infection (Angka et al., 2020). Such an approach could be key in treating severe cases of COVID-19, which correlate with cytokine storm and hyper-activation of immune responses normally not related to viral infections, particularly type-2 effectors (Lucas et al 2020). However, indiscriminate tempering of immune signaling would be counterproductive in the early stages of the infection and for patients with moderate disease, who appear to maintain a functional, well-adapted immune response.

DC/L-SIGN antagonists may help reducing the severity of SARS-CoV-2 infection by inhibiting the trans-receptor role played by these CLRs. We show here that Polyman26 (PM26), a glycomimetic antagonist of DC-SIGN, inhibits the interaction of the S protein with the lectin receptor and blocks DC-SIGN-mediated SARS-CoV-2 trans-infection of susceptible Vero cells. PM26 is known to act both by binding DC-SIGN CRD and by promoting its internalization (Berzi et al 2016), thus reducing the lectin concentration on cell surface and further impairing the ability of the virus to exploit it for enhancing the infection process. Additionally, upon binding to DC-SIGN PM26 was shown to induce a pro-inflammatory anti-viral response (Berzi et al., 2016), which should be beneficial at the onset of the infection and may help to steer the immune response towards a profile correlated with milder forms of the disease.

Demonstration of involvement of APCs in SARS-CoV-2 early dissemination through CRLs DC/L-SIGN, opens new avenues for understanding and treating the imbalanced innate immune response observed in COVID-19 pathogenesis. The potential of this and other CLRs antagonists in prevention or as part of combination therapy of COVID-19 needs to be further explored.

## Methods

### Protein production and purification

The extracellular domains (ECD) of DC-SIGN (residues: 66-404), L-SIGN (residues: 78-399), Langerin (residues: 68-328) and MGL (residues: 61-292) were produced and purified as already described (Achilli et al., 2020; Chabrol et al., 2012; Maalej et al., 2019; Reina et al., 2008) while SARS-CoV-2 Spike protein was expressed and purified as follows. The mammalian expression vectors used for the S ectodomain, derived from a pαH vector, was a kind gift from J. McLellan (Wrapp et al., 2020). This construct possesses, in its C-terminus, an 8xHis tag followed by 2 StrepTagII. EXPI293 cells grown in EXPI293 expression medium were transiently transfected with the S ectodomain vector according to the manufacturer’s protocol (Thermo Fisher Scientific). Cultures were harvested five days after transfection and the medium was separated from the cells by centrifugation. Supernatant was passed through a 0.45 µm filter and used for a two-step protein purification on Aktä Xpress, with a HisTrap HP column (GE Healthcare) and a Superose 6 column (GE Healthcare). Before sample loading, columns were equilibrated into 20 mM Tris pH 7.4; supplemented with variable concentrations of NaCl (150-500 mM) depending on the experiments. Unbound proteins were eluted from affinity column with equilibration buffer, contaminants with the same buffer supplemented with 75 mM imidazole while the spike protein was eluted with equilibration buffer supplemented with 500 mM imidazole and immediately loaded onto a gel filtration column run in equilibration buffer. Fractions of interest were pooled and concentrated at 0.5 mg/mL on an Amicon Ultra 50K centrifugal filter according to the manufacturer’s protocol (Millipore). The concentration of purified spike protein was estimated using an absorption coefficient (A_1%,1cm_) at 280 nm of 10.4 calculated using the PROTPARAM program (http://web.expasy.org/protparam/) on the Expasy Server. Quality control and visualization of the three different samples was performed by negative staining Transmission Electron Microscopy (TEM) using Uranyl Acetate as stain (2% w/v).

### Negative staining electron microscopy

Negative-stain grids were prepared using the mica-carbon flotation technique (Valentine et al., 1968). 4 µL of spike samples from purifications diluted at about 0.05-0.1 mg/mL were adsorbed on the clean side of a carbon film previously evaporated on mica and then stained using 2% w/v Uranyl Acetate for 30 s. The sample/carbon ensemble was then fished using an EM grid and air-dried. Images were acquired under low dose conditions (<30 e−/Å2) on a Tecnai 12 FEI electron microscope operated at 120 kV using a Gatan ORIUS SC1000 camera (Gatan, Inc., Pleasanton, CA) at 30,000x nominal magnification. To facilitate the visualization of the molecules, a Gaussian filter was applied to the images using Photoshop, then the gray levels were saturated and the background eliminated. For the 2D classification, images were processed with RELION 2.1 (Scheres, 2012). CTF was estimated with CTFFind-4.1 (Zhang, 2016). An initial set of 409 particles (box size of 512 pixels, sampling of 2.2 Å/pixel) was obtained by manual picking. After 2D classification the best looking 2D class averages were used as references for Autopicking. A set of around 35 000 particles (box size of 256 pixels, sampling of 4.4 Å/pixel, mask diameter 300Å) was obtained by Autopicking with a gaussian blob. The 8 best obtained classes were calculated from 2854 particles.

### SPR binding studies

Two types of surface plasmon resonance (SPR) experiments were performed at 25 °C on a Biacore T200. The first experiments with non-oriented spike surfaces were performed using a CM5 sensor chip, functionalized at 5 μL/min. Spike protein was immobilized on flow cells using a classical amine-coupling method. Flow cell 1 (Fc1) was functionalized with BSA and used as reference surface. Fc1 to 4 were activated with 50 μL of a 0.2 M EDC/ 0.05 M NHS mixture and functionalized with 20 µg/mL BSA (Fc1) or 50 μg/mL spike protein (Fc2-4), the remaining activated groups of all cells were blocked with 30 μL of 1 M ethanolamine pH 8.5. The four Fc were treated at 100 µL/min with 5 μL of 10 mM HCl to remove non-specificly bound protein and 5 μL of 50 mM NaOH/ 1M NaCl to expose surface to regeneration protocol. Finally, an average of 2500, 3000 and 2300 RU of spike protein were functionalized onto Fc2, 3 and 4, respectively.

The second type of experiments used oriented spike surfaces and were performed using a CM3 sensor chip functionalized at 5 μL/min. The procedure for oriented functionalization has been described in our recent work (Porkolab et al., 2020). Fc4 was functionalized with non-oriented spike protein exactly as described for the CM5 sensor chip using 20 µg/mL spike protein (final functionalization of 1430 RU). Fc1 (reference surface) and Fc2 were activated with 50 μL of a 0.2M EDC/ 0.05 M NHS mixture and functionalized with 170 μg/mL StrepTactin (IBA company) and the remaining activated groups were blocked with 80 μL of 1 M ethanolamine. Flow cells were treated at 100 µL/min with 5 μL of 10 mM HCl and 5 μL of 50 mM NaOH/ 1M NaCl. An average of 2000 RU of covalently immobilized StrepTactin was obtained. The spike protein was then captured at 20 µg/mL on Fc2 via its StrepTags. The surface was washed at 100 µL/min with 1M NaCl. Fc2 final level of functionalization was 990 RU. The presence of a double StrepTagII at the C-terminal extremity of the S protein (a total of 6 within the spike trimer) led to very stable surfaces and capture did not require the additional EDC/NHS treatment previously reported for such oriented functionalization. On both type of surfaces, for direct interaction studies, increasing concentrations of extracellular domain of DC-SIGN, L-SIGN, MGL and Langerin were prepared in a running buffer composed of 25 mM Tris pH 8, 150 mM NaCl, 4 mM CaCl_2_, 0.05% P20 surfactant, and either 80 μL of each DC-SIGN/L-SIGN ECD sample or 120 µL of Langerin/MGL ECD sample were injected onto the surfaces at 20 μL/min flow rate. The resulting sensorgrams were reference surface corrected. The apparent affinity of compounds was determined by fitting the steady state affinity model to the plots of binding responses versus concentration.

### Cell lines

Baby hamster kidney cells (BHK-21, 12-14-17 MAW, Kerafast, Boston, MA) and African Green Monkey Cell Line (VeroE6) were cultured in Dulbecco’s modified Eagle medium (DMEM) supplemented with 10% heat-inactivated fetal bovine serum (FBS), 25 μg/mL gentamycin and 2 mM L-glutamine. Jurkat, Jurkat DC-SIGN, Jurkat L-SIGN (Alvarez et al., 2002) and Jurkat langerin were maintained in RPMI 1640 supplemented with 10% heat-inactivated FBS, 25 μg/mL gentamycin and 2 mM L-glutamine.

### Production of human monocyte-derived macrophages and dendritic cells

Blood samples were obtained from healthy human donors (Hospital 12 de Octubre, Madrid, Spain) under informed consent and IRB approval. Peripheral blood mononuclear cells (PBMCs) were isolated by Ficoll density gradient centrifugation (Ficoll Paque, GE17-5442-02, Sigma-Aldrich). To generate monocyte-derived dendritic cells (MDDCs), 1 ×10^7^ PBMC/mL were placed into 24-well plates and incubated for 1 h at 37°C with 5% CO_2_. The adherent monocytes were then washed twice with PBS and resuspended in RPMI supplemented with the cytokines GM-CSF (1000 U/mL) and IL-4 (500 U/mL) (Miltenyi Biotec). Differentiation to immature MDDCs was achieved by incubation at 37 °C with 5% CO_2_ for 7 days and subsequent activation with cytokines addition every other day. To generate monocyte-derived macrophages (M2-MDMs), CD14+ monocytes were purified using anti-human CD14 antibody-labeled magnetic beads and iron-based LS columns (Miltenyi Biotec) and used directly for further differentiation into macrophages (Dominguez-Soto et al., J Immunol 2011). For differentiation of M2-MDMs, cells were incubated at 37°C with 5% CO_2_ for 7 days and activated with M-CSF (1000 U/mL) (Miltenyi Biotec) every second day.

### Production of SARS-CoV-2 pseudotyped recombinant Vesicular Stomatitis Virus (rVSV-luc)

rVSV-luc pseudotypes were generated following a published protocol (Whitt, J Virol Methods 2010). First, BHK-21 were transfected to express the S protein of SARS-CoV-2 (codon optimized, kindly provided by J. García-Arriaza, CNB-CSIC, Spain), Ebola virus Makona Glycoprotein (EBOV-GP) (KM233102.1) was synthesized and cloned into pcDNA3.1 by GeneArt AG technology (Life Technologies, Regensburg, Germany) or VSV-G following the manufacturer’s instructions of Lipofectamine 3000 (Fisher Scientific). After 24 h, transfected cells were inoculated with a replication-deficient rVSV-luc pseudotype (MOI: 3-5) that contains firefly luciferase instead of the VSV-G open reading frame, rVSVΔG-luciferase (G*ΔG-luciferase) (Kerafast, Boston, MA). After 1 h incubation at 37°C, cells were washed exhaustively with PBS and then DMEM supplemented with 5% heat-inactivated FBS, 25 μg/mL gentamycin and 2 mM L-glutamine were added. Pseudotyped particles were collected 20-24 h post-inoculation, clarified from cellular debris by centrifugation and stored at −80°C (Hoffmann et al., 2020; Letko et al., 2020; Whitt, 2010). Infectious titers were estimated as tissue culture infectious dose per mL by limiting dilution of rVSV-luc-pseudotypes on Vero E6 cells. Luciferase activity was determined by luciferase assay (Steady-Glo Luciferase Assay System, Promega).

### Direct infection

Cell lines: Jurkat, Jurkat DC-SIGN and Jurkat L-SIGN (3 × 10^5^ cells) or primary cells: MDDCs, M2-MDMs (5 × 10^4^ cells) were challenged with SARS-CoV-2, EBOV-GP or VSV-G pseudotyped recombinant viruses (MOI: 0.5-2). After 24 h of incubation, cells were washed twice with PBS and lysed for luciferase assay.

### Trans-infection

For trans-infection studies, Jurkat DC-SIGN, Jurkat L-SIGN, Jurkat Langerin (3 × 10^5^ cells) or MDDCs (5 × 10^4^ cells) were challenged with recombinant SARS-CoV-2, EBOV-GP or VSV-G pseudotyped viruses (MOI: 0.5-2) and incubated during 2 h at room temperature with rotation. Cells were then centrifuged at 1200 rpm for 5 minutes and washed twice with PBS supplemented with 0.5% bovine serum albumin (BSA) and 1 mM CaCl_2_. Jurkat DC-SIGN and MDDCs were then resuspended in RPMI medium and co-cultivated with adherent Vero E6 cells (1.5 × 10^5^ cells/well) on a 24-well plate. After 24 h, the supernatant was removed and the monolayer of Vero E6 was washed with PBS three times and lysed for luciferase assay.

### Synthesis of Inhibitors and inhibition experiments

Polyman26 (PM26) is a known glycomimetic ligand of DC-SIGN and an antagonist of DC-SIGN mediated HIV trans-infection (Berzi et al., 2016; Ordanini et al., 2015). It was synthesized as previously described and tested in SPR studies as an inhibitor of DC-SIGN interaction to the spike protein of SARS-CoV-2, using both the oriented and non-oriented S surface described above. In both cases, a 20 µM solution of DC-SIGN in a running buffer composed of 25 mM Tris pH 8, 150 mM NaCl, 4 mM CaCl_2_, 0.05% P20 surfactant was co-injected with variable concentrations of Polyman26, from 50 µM to 0.1 µM, in the same buffer. IC_50_ values were determined from the plot of PM26 concentration vs % inhibition by fitting four-parameter logistic model as previously described (Varga et al., 2014).

In the infection studies, cells were first incubated with the compound PM26 for 20 min at room temperature with rotation and then challenged with SARS-CoV-2 recombinant viruses (MOI: 0.5-2) during 2 h at room temperature with rotation. The concentrations tested for compound PM26 were 5 and 0.5μM. As a control, inhibition experiment was performed in the presence of anti-DC/L-SIGN antibody (R&D Systems). Cells were then washed as described above, resuspended in RPMI medium and co-cultivated with adherent Vero E6 cells (1.5 × 10^5^ cells/well) on a 24-well plate. After 24 h, the supernatant was removed and the monolayer of Vero E6 was washed with PBS three times and lysed for luciferase assay.

## Acknowledgments

This work used the platforms of the Grenoble Instruct-ERIC centre (ISBG; UMS 3518 CNRS-CEA-UGA-EMBL) within the Grenoble Partnership for Structural Biology (PSB), supported by FRISBI (ANR-10-INBS-05-02) and GRAL, within the University Grenoble Alpes graduate school CBH-EUR-GS (ANR-17-EURE-0003). The EM facility is supported by the Auvergne-Rhône-Alpes Region, the Fondation Recherche Medicale, the fonds FEDER and the GIS-Infrastructures Biologie Sante et Agronomie (IBISA). F.F. acknowledges the French Agence Nationale de la Recherche PIA for Glyco@Alps (ANR-15-IDEX-02). Research in R.D. lab is supported by grants from the Instituto de Investigación Carlos III, ISCIII, (FIS PI 1801007), the European Commission Horizon 2020 Framework Programme: Project VIRUSCAN FETPROACT-2016: 731868, and by Fundación Caixa-Health Research (Project StopEbola). F.F. thanks J. McLellan for sharing expressing vector and E. Fadda for making models of the spike glycoprotein available as well as for stimulating exchanges on the #glycotime on twitter.

## Author contributions

R.D. and F.F. conceived, designed, supervised the work and wrote first draft of the manuscript. A.B. contributed to the writing, M.T., J.L., C.V., N.L., I.B., F.L., Y.G., S.S., N.T. performed the experiment, D.F., G.S., processed EM Data, S.S., A.B. synthesized glycomimetic compound, R.D., F.F., S.S., A.B., N.T and G.S. reviewed and edited the manuscript. All authors have read and approved the final manuscript.

