## Supplementary material for "DC/L-SIGN recognition of spike glycoprotein promotes SARS-CoV-2 trans-infection and can be inhibited by a glycomimetic antagonist": supp. info

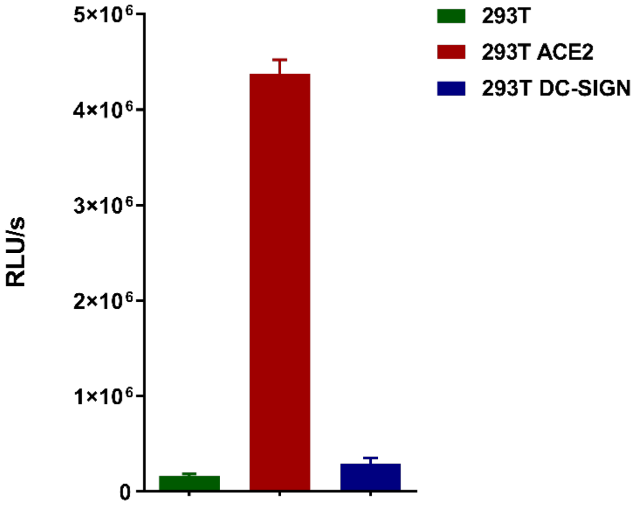


**Fig. S1. Infection of 293T-ACE2 and 293T-DC-SIGN with SARS-CoV-2**. 293T were transfected to express ACE2 or DC-SIGN on the cell surface. Parental non-permissive 293T were used as a negative control of infectivity. Results are represented as bars corresponding to mean±SEM of infection with VSV/SARS-COV-2 pseudovirions measured by triplicates. Infection values from luciferase assays are expressed as Relative Light Units (RLUs).

#### DC-SIGN does not function as an alternative receptor for SARS-CoV-2

The potential function of DC-SIGN as an alternative receptor for SARS-CoV-2 was also evaluated in 293T cell line transfected to express DC-SIGN. Parental 293T are not susceptible to SARS-CoV-2 infection, thus they were used as a negative control of infection. On the other hand, 293T expressing ACE2 were used a positive control of SARS-CoV-2 infection.

Parental 293T lack ACE2 expression and as expected were not infected by VSV/SARS-COV-2 pseudovirions. The expression of ACE2 on 293T was sufficient for efficient infection with SARS-COV-2 (Fig. S1). Nevertheless, the expression of DC-SIGN on 293T surface did not allow SARS-COV-2 infection, indicating that DC-SIGN does not function as alternative receptor to ACE2 in non-permissive cells.

### Methods

#### Cell lines (add to the part of adherent cell lines)

Human embryonic kidney cells (293T; ATCC-CRL-11268) were cultured in DMEM supplemented with 10% heat-inactivated FBS, 25 µg/mL gentamycin and 2 mM L-glutamine.

**Infection of 293T-ACE2 and 293T-DC-SIGN with SARS-CoV-2**

293T cells were plated in a 24-well tissue culture plate ($5 \times 10^5$ cells/well) and the next day cells were transfected with phACE2 (kindly provided by Andrea Cara, Instituto Superiore di Sanità, Rome, Italy) or with pLZR-DC-SIGN-CITE-GFP (Alvarez et al., J Virol 2002) using a standard Lipofectamine 3000 transfection protocol (Fisher Scientific). After 24h, transfected cells were challenged with VSV/SARS-CoV-2 (MOI: 2). Following 24h of incubation cells were washed twice with PBS and lysed for luciferase assay (Steady-Glo Luciferase Assay System, Promega). Parental 293T were used as a negative control of infection.
